# Changes in local interaction rules during ontogeny underlie the evolution of collective behavior

**DOI:** 10.1101/2023.03.28.534467

**Authors:** Alexandra Paz, Karla J. Holt, Anik Clarke, Ari Aviles, Briana Abraham, Alex C. Keene, Erik R. Duboué, Yaouen Fily, Johanna E. Kowalko

## Abstract

Collective motion emerges from individual interactions which produce groupwide patterns in behavior. While adaptive changes to collective motion are observed across animal species, how local interactions change when these collective behaviors evolve is poorly understood. Here, we use the Mexican tetra, *A. mexicanus,* which exists as a schooling surface form and a non-schooling cave form, to study differences in how fish alter their swimming in response to neighbors across ontogeny and between evolutionarily diverged populations. We find that surface fish undergo a transition to schooling during development that occurs through increases in inter-individual alignment and attraction mediated by changes in the way fish modulate speed and turning relative to neighbors. Cavefish, which have evolved loss of schooling, exhibit neither of these schooling-promoting interactions at any stage of development. These results reveal how evolution alters local interaction rules to produce striking differences in collective behavior.

## Introduction

Social behaviors in animals are critical for survival, and extensive variation in sociality is found across animal species. Collective motion, which includes flocking in birds, herd migration in ungulates, and swarming in insects, is an example of collective behavior in which individuals’ responses to local social cues culminate in coordinated behavioral outcomes (Ahmed & Faruque, 2022; Bialek et al., 2012; Fullman et al., 2021; Götmark et al., 1986; Ling et al., 2019; Nagy et al., 2010; Naidoo et al., 2012; Wang et al., 2021). Collective motion is also observed in many species of fish, and includes shoaling and schooling (Greenwood et al., 2013; Ho et al., 2015; Katz et al., 2011; Seghers, 1974; Suriyampola et al., 2016; Z.-H. Tang et al., 2017). Shoaling is defined as fish maintaining close proximities to other individuals in the group, while schooling is characterized by fish maintaining both close proximity and alignment. While shoaling and schooling result in complex collective motion in groups of up to thousands of individuals, these group dynamics emerge from local interactions between individuals within the group, such as individuals moving toward or away from neighbors based on their relative position (Bierbach et al., 2020; Herbert-Read et al., 2011, 2017; Katz et al., 2011). How these local interactions manifest in group level dynamics has been established in various fish species that display robust schooling and shoaling (Bierbach et al., 2020; Harpaz et al., 2021; Herbert-Read et al., 2019; Hinz & de Polavieja, 2017; Ioannou et al., 2017; Jolles et al., 2017; Katz et al., 2011; Tunstrøm et al., 2013). However, how evolution impacts these local interaction rules to produce group level differences is not understood. Establishing how changes to individual behaviors lead to variation in collective motion is critical to revealing how collective behaviors evolve in natural populations.

The Mexican tetra, *Astyanax mexicanus,* is a species of freshwater fish that consists of surface populations which inhabit rivers and streams in Mexico and Southern Texas, and multiple independently evolved cave fish populations that inhabit caves in Northeastern Mexico (Espinasa et al., 2018; Herman et al., 2018; Jeffery, 2009; Mitchell et al., 1977). Caves inhabited by *A. mexicanus* have a number of differences in ecology relative to the surface habitat, including constant darkness, loss of macroscopic predators and differences in water chemistry (Boggs & Gross, 2021; Elliott, 2018; Fish, 1977; Mitchell et al., 1977; Ornelas-García et al., 2018; Rohner et al., 2013; Tabin et al., 2018). These ecological differences have resulted in the repeated evolution of a number of morphological, physiological and behavioral traits in *A. mexicanus* cave fish relative to their surface conspecifics, including loss or reduction of eyes and pigmentation, enhancement of non-visual sensory systems, changes to metabolism and reductions in sleep (Alié et al., 2018; Bibliowicz et al., 2013; Borowsky, 2016; Chin et al., 2018; Duboué et al., 2011; Jeffery, 2009; Klaassen et al., 2018; Lloyd et al., 2018; Protas & Jeffery, 2012; Yoffe et al., 2020; Yoshizawa et al., 2014). *A. mexicanus* cave fish have also evolved changes to multiple social behaviors relative to surface fish, including reduced aggression and an absence of social hierarchies (Breder, 1943; Burchards et al., 1985; Elipot et al., 2013; Espinasa et al., 2022; Langecker et al., 1995). Further, while adult surface fish exhibit robust shoaling and schooling in the lab and in the field, these behaviors are reduced in adult fish from multiple cave fish populations (Gregson & Burt de Perera, 2007; Iwashita & Yoshizawa, 2021; John, 1964; Kowalko et al., 2013; Patch et al., 2022). Thus, the robust differences in schooling and shoaling between surface and cave fish provide an opportunity to investigate how individual interactions are altered over evolutionary time to produce differences in group behaviors.

Here, we quantify group dynamics and individual behaviors in groups of cave and surface *A. mexicanus* across ontogeny to identify how changes in individual fish behaviors lead to evolutionary loss of collective motion. By examining inter-individual interactions across development, we are able to identify when during development fish initially begin to modulate their motion relative to their neighbors, and how these changes in individual behaviors alter group level behaviors. Through comparing inter-individual interactions across populations that exhibit markedly different group level behaviors, we define how different social interaction rules underlying group behaviors have evolved, as well as when the developmental trajectories leading to different group level behaviors diverge.

## Results

### Attraction and alignment diverge between surface and cave fish over the course of development

The ontogeny of schooling and shoaling in surface fish, as well as the stage at which cave and surface fish social behaviors diverge, is unknown. For example, surface and cave fish may display distinct social interactions throughout development, or alternatively, they may initially display similar interactions before diverging later in development. To determine if schooling and shoaling change across development, we analyzed swimming behavior in surface fish and cave fish in groups of five at timepoints across development: 7 days post fertilization (dpf), shortly after fish begin to hunt prey and feed, 28 dpf, 42 dpf, and 70 dpf, when fish have reached subadult stages, but prior to sexual maturity. We calculated the distance (fig 1a) and the alignment (fig 1b) between pairs of fish, as these metrics have previously been used to define schooling and shoaling (Jolles et al., 2017; Katz et al., 2011; Patch et al., 2022; W. Tang et al., 2020) (fig1c). The joint probability distributions of pair distance and pair angle spread over the entire range of distances and angles in surface *A. mexicanus* at larval and juvenile time points (7 and 28 dpf; fig 1d, e), suggesting that surface fish at these stages of development are neither schooling nor shoaling. However, while the pair distances and angles continue to spread across the entire range at 42 dpf, by this point in development the distribution also exhibits a peak at short pair distance and small pair angle, suggesting some preference for proximity and alignment by this stage (fig 1f). In groups of 70 dpf surface fish, the joint probability distribution of pair distance and angle has a sharp peak at short pair distances, which is more pronounced at small pair angles, suggesting that surface fish show a strong preference for proximity and alignment at this stage (fig 1g). These 70 dpf results indicate that fish are schooling and shoaling at this stage. Together these data suggest that in surface fish, schooling and shoaling emerge over the course of development, with an initial preference for being both aligned and in close proximity beginning prior to 42 dpf and becoming robust by 70 dpf.

**Figure 1.**
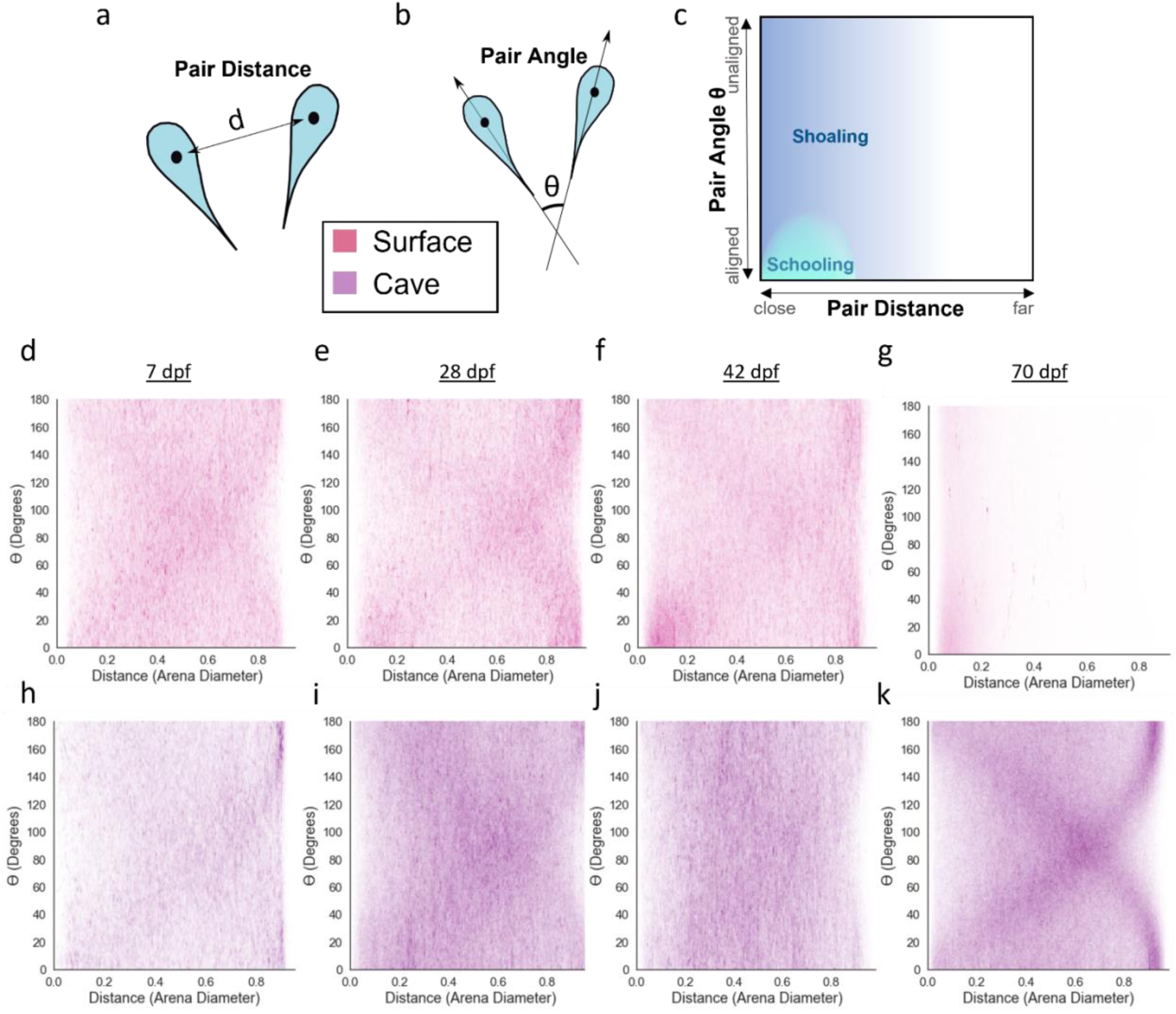
Joint probability distributions of pair distance and angle between individuals in groups of five fish. **a)** Pair distance was defined as the distance between the center points of 2 individuals. **b)** Pair angle was defined as the difference in heading between 2 individuals. **c)** Collective behaviors such as schooling and shoaling can be broadly defined using the relationship between pair proximity and orientation. Joint plots for groups of surface fish at **d)** 7 dpf, **e)** 28 dpf, **f)** 42 dpf, and **g)** 70 dpf. Joint plots for groups of cave fish at **h)** 7 dpf, **i)** 28 dpf, **j)** 42 dpf, or **k)** 70 dpf.

To determine if evolutionary loss of schooling in cavefish occurs at late developmental stages, or if surface and cave fish behavioral differences in sociality can be observed throughout developmental stages, we assessed whether cave fish demonstrate a preference for alignment and proximity at any point in development. Similar to surface fish at the same stages, the joint probability distributions of pair distance and pair angle for 7 dpf and 28 dpf cave fish groups spread over the entire available range of distances and angles (fig 1h, i). However, while surface fish begin to exhibit patterns of inter-fish proximity and alignment associated with schooling and shoaling at 42 dpf, cave fish do not. Instead, the joint probability distribution of pair distance and pair angle continues to spread over a range of distances and angles in 42 dpf and 70 dpf cave fish groups, suggesting a lack of preference for proximity or alignment at these later developmental stages (fig 1j, k). A different pattern emerges in 70 dpf cave fish: a pair of arches that suggests a strong preference for swimming along the arena walls (fig 1k) (Patch et al., 2022). These findings suggest that cave fish do not school or shoal at any point in development. Taken together, these results indicate that the attraction and alignment that underlie schooling behavior emerge in surface fish over the course of development, with attraction and/or alignment being present at 42 dpf, and that these behaviors do not follow this developmental trajectory in cavefish.

### Surface fish develop the tendency to align prior to attraction

We next asked if attraction and tendency to align to neighbors are established at the same developmental stages in surface fish. To determine when in development surface fish begin to exhibit a preference for alignment to one another, we compared the angles of fish to their nearest neighbors across ontogeny. Alignment of nearest neighbors was compared to alignment of nearest neighbors in mock groups generated by extracting the positions of individuals that were not assayed together and combining them to form groups of five fish (see methods). Comparison to mock groups allows us to account for factors that may differ over development, but that are not directly related to collective behavior, such as differences in locomotion unrelated to social behavior and tendency to align with the walls. At 7 dpf, surface fish nearest neighbor alignment was similar to mock groups (actual median = 85.3°, mock median = 88.1°; fig 2a), suggesting that at early stages of development, surface fish do not have a preference for alignment. Beginning at 28 dpf, however, there is a statistically significant decrease in pair angle in actual groups of surface fish compared to pair angle in mock groups (actual median = 81.6°, mock median = 88.4°; fig 2a), indicating that fish aligned more with their neighbors than expected by chance. At 42 dpf and 70 dpf, alignment between nearest neighbors relative to alignment in mock groups became more pronounced than at earlier developmental timepoints (42 dpf: actual median = 79.2°, mock median = 87.0; 70 dpf: actual median = 58.7°, mock median = 81.8°; fig 2a), suggesting that in surface fish, the tendency to align with nearest neighbors becomes more pronounced over the course of development. In contrast, the alignment of nearest neighbors in groups of cave fish did not significantly differ from the alignment of mock group nearest neighbor pairs at any of the developmental timepoints (7 dpf: actual median = 83.6°, mock median = 86.5°; 28 dpf: real median = 87.6°, mock median = 89.3°; 42 dpf: real median = 89.6°, mock median = 89.3°; 70 dpf: real median = 90.3°, mock median = 88.9°; fig 2b). Importantly, the nearest neighbor pair angles of surface and cave fish did not strongly correlate with swimming speed at any developmental stage, suggesting the observed trends are not simply the consequence of differences in swimming speed (Fig S1a & S1b). Together, these data suggest that a preference to align with neighbors emerges between 7 dpf and 28 dpf in surface fish and increases over time. By contrast, cave fish demonstrate no tendency to align with neighbors at any timepoint, suggesting that loss of tendency to align contributes to loss of schooling in cavefish.

**Figure 2.**
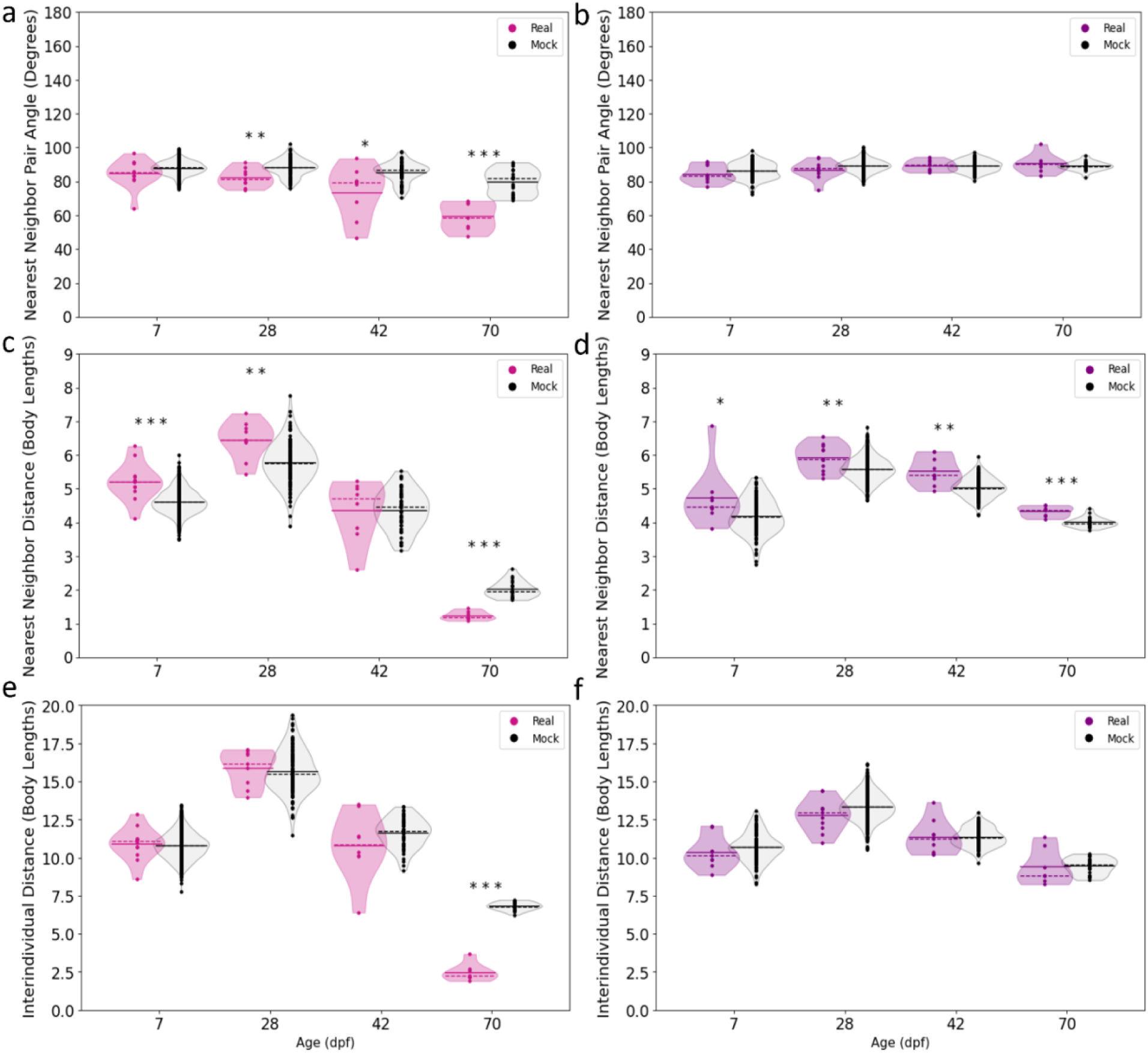
Comparisons of surface and cave fish proximity and alignment to that of mock groups. **a)** Comparisons of surface fish (left violin plots, pink) nearest neighbor pair angles to mock groups (right violin plots, gray) at 28 dpf (Real x̃ = 81.6, Mock x̃ = 88.5, U = 243.0, p = 0.004), 42 dpf (Real x̃ = 81.6, Mock x̃ = 87.0, U = 107.0, p = 0.016), and 70 dpf (Real x̃ = 58.7, Mock x̃ 81.8, U = 0.0, p < 0.001) **b)** Comparisons of cave fish (left violin plots, purple) nearest neighbor pair angle to mock groups (right violin plots, gray) at 7 dpf (Real x̃ = 83.6, Mock x̃ = 86.5, U = 435.0, p = 0.246), 28 dpf (Real x̃ = 87.6, Mock x̃ = 89.3, U = 1855.0, p = 0.126), 42 dpf (Real x̃ = 89.6, Mock x̃ = 89.3, U = 574.0, p = 0.954), and 70 dpf (Real x̃ = 90.3, Mock x̃ = 88.9, U = 85.0, p = 0.568). **c)** Comparisons of surface fish nearest neighbor distances to mock groups at 7 dpf (Real x̃ = 5.21, Mock x̃ = 4.60, U = 7998.0, p < 0.001), 28 dpf (Real x̃ = 6.46, Mock x̃ = 5.75, U = 905.0, p = 0.003), 42 dpf (Real x̃ = 4.70, Mock x̃ = 4.56, U = 247.0, p = 0.654), and 70 dpf (Real x̃ = 1.18, Mock x̃ = 1.95, U = 0.0, p < 0.001). **d)** Nearest neighbor distances of cave fish compared to mock groups at 7 dpf (Real x̃ = 4.46, Mock x̃ = 4.16, U = 795.0, p = 0.045), 28 dpf (Real x̃ = 5.89, Mock x̃ = 5.58, U = 3729.0, p = 0.008), 42 dpf (Real x̃ = 5.40, Mock x̃ = 5.02, U = 929.0, p = 0.001), and 70 dpf (Real x̃ = 4.38, Mock x̃ = 3.97, U = 136.0, p < 0.001). **e)** Comparisons of surface fish interindividual distance to mock groups at 7 dpf (Real x̃ = 11.07, Mock x̃ = 10.79, U = 5303.0, p = 0.490), 28 dpf (Real x̃ = 16.13, Mock x̃ = 15.47, U = 638.0, p = 0.534), 42 dpf (Real x̃ = 10.84, Mock x̃ = 11.7, U = 167.0, p = 0.257), and 70 dpf (Real x̃ = 2.28, Mock x̃ = 6.79, U = 0.0, p < 0.001). **f)** Interindividual distances in groups of cave fish compared to mock groups at 7 dpf (Real x̃ = 10.13, Mock x̃ = 10.67, U = 408.0, p = 0.162), 28 dpf (Real x̃ = 12.95, Mock x̃ = 13.32, U = 1784, p = 0.091), 42 dpf (Real x̃ = 11.25, Mock x̃ = 11.30, U = 505.0, p = 0.587), and 70 dpf (Real x̃ = 8.83, Mock x̃ = 9.52, U = 60.0, p = 0.499). Solid lines denote means, dotted lines denote medians, each point denotes a single trial, * denotes p < 0.05, ** denotes p < 0.01, *** p < 0.001. Real = real data, Mock = mock data, x̃ = median, α = 0.05.

In addition to alignment, attraction to others in the group is essential for schooling and shoaling behavior in fish. To determine when fish first begin to show attraction to other fish in the group, we calculated two commonly used metrics of attraction: nearest neighbor distance, the distance between each fish and its closest neighbor, and interindividual distance, the distances between all pairs of fish in the group, a measure of group coherency. At 7 dpf and 28 dpf, surface fish nearest neighbor distances were significantly larger than those of mock groups (fig 2c; 7 dpf real NND median = 5.2 body lengths (BLs), mock NND median = 4.6 BLs; 28 dpf real NND median = 6.5 BLs, mock NND median = 5.8 BLs). This changes at 42 dpf, when surface fish nearest neighbor distances were similar between real and mock groups (real NND median = 4.8 BL, mock NND median = 4.5 BL; fig 2c). By 70 dpf, however, surface fish maintained significantly closer nearest neighbor distances compared to mock groups (real median = 1.2 BLs; mock median = 2.0 BLs; fig 2c). These data suggest that surface fish do not display attraction during early development, and may only avoid neighbors at these developmental stages. Further, they suggest that surface fish begin to display robust attraction by 70 dpf. This establishment of attraction by 70 dpf is also observed at the level of group coherency. At 7, 28, and 42 dpf, surface fish interindividual distances resembled those of mock groups (7 dpf real IID median = 11.1 BLs, mock IID median = 10.8 BLs; 28 dpf real IID median = 16.2 BLs, mock IID median = 15.5 BLs; 42 dpf real IID median = 10.8 BLs, mock IID median = 11.7 BLs; fig 2e). However, at 70 dpf, surface fish groups show cohesiveness, with interindividual distances that are significantly smaller compared to mock groups (real median = 2.3 BLs; mock median = 6.8 BLs; fig 2e).

Across development, cave fish maintained significantly greater distances from their nearest neighbors compared to mock groups (7 dpf: real NND median = 4.5 BLs, mock NND median = 4.2 BL; 28 dpf: real NND median = 5.9 BLs, mock NND median = 5.6 BLs; 42 dpf: real NND median = 5.4, mock NND median = 5.0 BLs; 70 dpf: real NND median = 4.4 BLs, mock NND median = 4.0 BLs; fig 2d). However, the interindividual distances in groups of cave fish resembled those of control mock groups over the course of all developmental timepoints assayed (7 dpf: real IID median = 10.1 BLs, mock IID median = 10.7 BLs; 28 dpf: real IID median = 13.0 BLs, mock IID median = 13.3 BLs; 42 dpf: real IID median = 11.2 BLs, mock IID median = 11.3 BLs; 70 dpf: real IID median = 8.8 BLs, mock IID median = 9.5 BLs; fig 2f). These data suggest that cave fish do not exhibit attraction to neighbors at any point in development, and may exhibit repulsion from nearest neighbors. We found no strong correlations between swimming speed and nearest neighbor distance or interindividual distance in surface fish or cave fish at most stages, except at 70 dpf where it correlated with approximately 20% of the variability observed in interindividual distance in cave fish, suggesting the observed trends are not simply the consequence of differences in swimming speed (Fig S1a & S1b). Taken together, these data suggest that surface fish attraction and preference for alignment develop at distinct points in development, and that lack of a preference for alignment or attraction is maintained in cavefish across ontogeny.

### Reductions in tendency to modulate speed and turning according to neighbor position underlie loss of schooling in cave fish

We next sought to understand how fish modulate the ways they interact with their neighbors that give rise to the emergence of schooling and shoaling, and how these inter-fish interactions differ between populations that have evolved differences in the tendency to school and shoal. We first defined inter-fish positional preferences by generating density heat maps around a focal fish and looking for differences between real groups of fish and mock groups. At 70 dpf, when surface fish school and shoal, they exhibit specific positional preferences relative to neighboring fish when compared to mock groups: Fish are frequently positioned such that neighboring fish occupy a zone between 0.05 and 0.4 tank radii from a focal fish, a zone we refer to as the schooling zone (fig 3a, fig S2a). In contrast, fish in 70 dpf surface fish mock groups do not preferentially occupy the schooling zone (fig S3a). This positional preference is also observed, to a lesser extent, at 42 dpf (fig 3a). However, it is not present at earlier developmental stages. Instead, density maps of 7 and 28 dpf surface fish indicate low fish density at close distances relative to the focal fish (fig 3a). This lack of preference for proximity is also observed across development in cave fish (fig 3b). Together, these data suggest that neither surface nor cave fish display robust attraction early in development, however surface fish develop attraction over the course of development, consistent with the emergence of schooling behavior in these fish.

**Figure 3:**
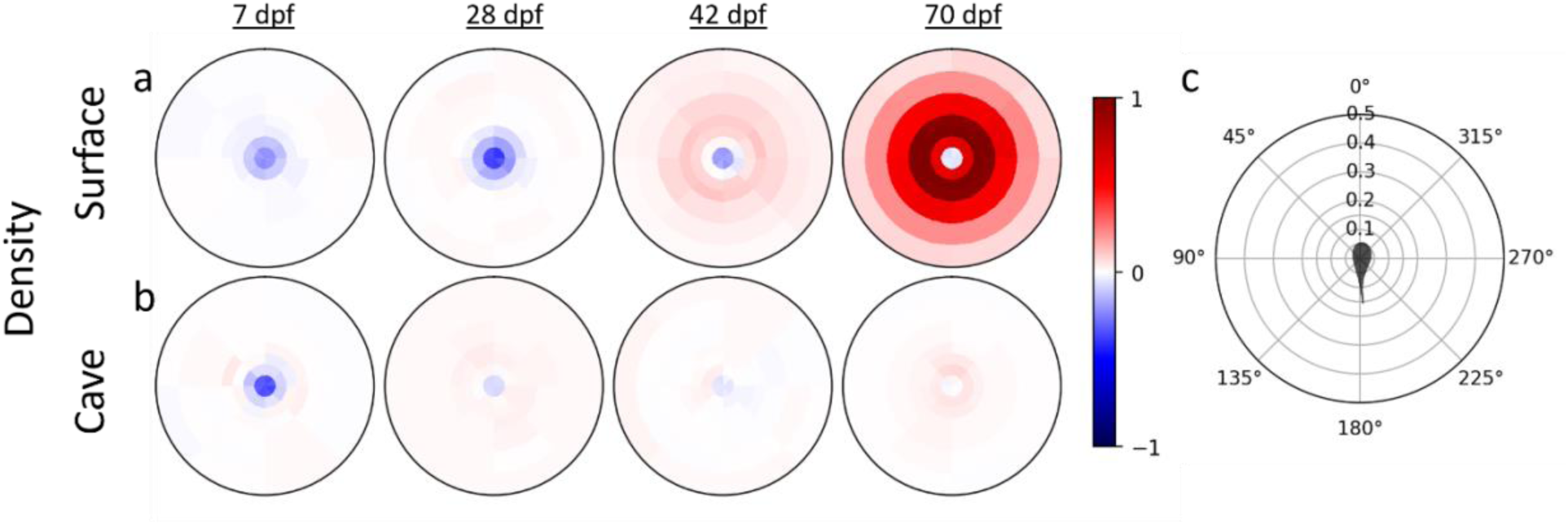
Positional preferences of surface and cave fish across development compared to mock groups. Density heat maps illustrate the preferred positions of **a)** surface and **b)** cave fish individuals relative to a focal fish located at the center of the heat map, facing upwards. Values close to 1 (red) indicate high fish density and values close to −1 (blue) indicate very low fish density. Results here are the difference between real (fig S2) and mock groups of 5 fish (fig S3). c) Polar grid. The focal fish is shown in the center, facing up. The rings at 0.05, 0.1, 0.15, 0.2, 0.3, 0.4, and 0.5 tank radii show the distance between the focal fish and its neighbor.

Fish can modulate their position relative to neighbors using a combination of two behaviors: turning and changing swimming speed, and one or both of these behaviors could be altered by evolution in cave fish. In order to determine the contributions of speed changes and turning to the maintenance of preferred positions, we computed the average speeding and turning forces of each fish when another fish is nearby as a function of the neighboring fish’s location, similar to previous work in golden shiners (Katz et al., 2011). Force here refers to the focal fish’s acceleration normalized to average speed. The turning force is the normal acceleration (acceleration perpendicular to the fish’s heading). The speeding force is the tangential acceleration (acceleration in the direction of the fish’s heading). In order to control for the effects of arena walls, speeding and turning force were also calculated for individuals in mock groups, and the difference between real (fig S2) and mock group data (fig S3) were plotted as heat maps (fig 4ab). At 70 dpf, surface fish tend to increase swimming speed if a neighbor is located further than ∼0.15 tank radii in front of them and decrease swimming speed if a neighbor is located further than ∼0.15 tank radii behind them (fig 4a). Since 0.15 tank radii corresponds to the peak of the neighbor density heatmap (fig 3a), this suggests that 0.15 tank radii is the preferred distance between schooling neighbors, and that fish modulate their speed to get closer to their neighbor when that neighbor is further away than this preferred distance.

**Figure 4:**
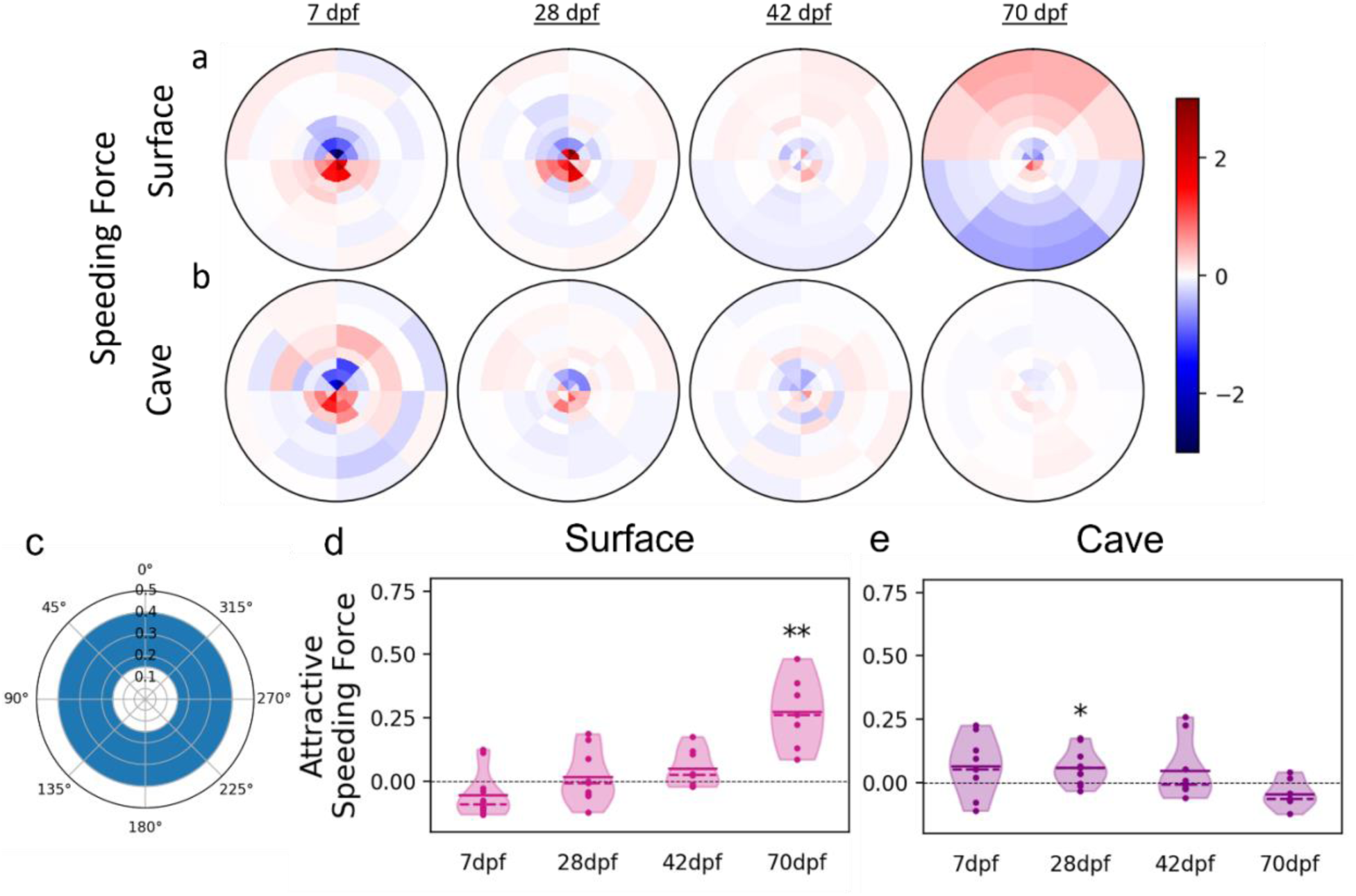
Fish modulate swimming speed according to the relative position of neighbors. Speeding force represents the acceleration of an individual **a)** surface or **b)** cave fish in the axis of its motion. Speeding force here is given as a function of the position of neighboring fish, with the focal fish’s direction of motion oriented toward the top of the figure. Values close to 1 (red) indicate increases in swimming speed and values close to −1 (blue) indicate decreases in swimming speed. **c)** The mean speeding force for each trial was calculated for pairs within proximities in which attraction was expected (0.15 and 0.4 tank radii) to give the attractive speeding force. **d)** The mean attractive speeding forces of surface fish at 7 dpf (x̃ = −0.055, t = 20, p = 0.151), 28 dpf (x̃ = 0.0189, T = 0.536, p = 0.606), and 42 dpf (x̃ = 0.051, T = 1.96, p = 0.091), and 70 dpf (x̃ = 0.274, T = 5.14, p = 0.002)**. e)** The mean attractive speeding forces of cave fish at 7 dpf (x̃ = 0.067, T = 1.77, p = 0.115), 28 dpf (x̃ = 0.061, T = 2.97, p = 0.014), 42 dpf (x̃ = 0.047, T = 20, p = 0.820), and 70 dpf (x̃ = −0.043, T = −2.01, p = 0.091). Accelerations here have been standardized to mean swimming speed of each age and population. ** denotes p < 0.01, x̃ = mean, α = 0.05. Results here are the difference between real and mock data.

In order to quantitatively assess the contribution of speed changes to attraction, we calculated the mean attractive speeding force for all pairs of fish within the attraction zone, defined as the region between 0.15 and 0.4 tank radii of a focal fish, i.e., the part of the schooling zone where the speeding force is expected to be attractive (fig 4c). Positive values indicate that speed changes tend to decrease the distance between neighbors whereas negative values indicate that speed changes tend to increase the distance between neighbors. Values close to zero indicate that individuals are not utilizing changes in swimming speed to change their distance relative to neighbors. To account for speeding due to non-social effects, we subtracted the mock group mean from the mean of each real group. At 70 dpf, the mean trial speeding force of surface fish is significantly greater than zero (mean = 0.274 cm/s^2^, p = 0.002), indicating that fish show a tendency to use speed to position themselves closer to neighbors at this stage (fig 4d). At 7, 28 and 42 dpf, surface fish do not appear to modulate speed in response to neighbors in the attraction zone, and mean trial speeding forces do not differ significantly from zero (7 dpf mean = −0.055 cm/s^2^, p = 0.151; 28 dpf mean = 0.0189 cm/s^2^, p = 0.606; 42 dpf mean = 0.051 cm/s^2^, p = 0.091) (fig 4a & d).

Cave fish attractive speeding force did not significantly differ from zero at 7 dpf (mean = 0.067 cm/s^2^, p = 0.115) or 42 dpf (mean = 0.047 cm/s^2^, p = 0.820), although speeding force was slightly greater than zero at 28 dpf (mean = 0.061 cm/s^2^, p = 0.014) (fig 4e). Unlike in surface fish, cave fish speeding forces at 70 dpf (mean = −0.043 cm/s^2^, p = 0.091) also did not differ from zero, consistent with the positional preferences and observed lack of schooling and shoaling at these stages. Considering the similarities in surface and cave fish positional preferences at early stages (fig 3a & 3b), we hypothesized that surface and cave fish may exhibit similar trends in speeding force at close proximities during this stage. Within close proximities, surface fish at 7 and 28 dpf slow down when fish are in front of them and speed up when fish are behind them, a trend that continues across time points and is present in cave fish (fig 4a, b). Together, these data suggest that both surface and cave fish alter their speed to maintain positional preferences relative to neighbors, but only surface fish develop the tendency to modulate speed to get closer to neighbors, contributing to maintenance of close proximity required for schooling and shoaling.

Next, we assessed whether fish modulate their position relative to other fish through turning. Surface fish at 70 dpf turn toward neighbors located any greater than 0.15 tank radii to the right or left of them (fig 5a). Similar to speeding force, we calculated the attractive turning force (positive if the turn is towards neighbor, negative if it’s away from it) averaged over every pair of fish within the attraction zone (fig 5c). To account for turning due to non-social effects, we subtracted the mock group mean from the mean of each real group. At 70 dpf, the attractive turning force in surface fish is significantly higher than zero (mean = 0.270 *Rad*⁄*s*^2^, p = 0.002) (fig 5d). A slight tendency to turn toward neighbors can be observed in heat maps at 42 dpf, and an attractive turning force slightly higher than zero, though the difference was not statistically significant (mean = 0.071 *Rad*⁄*s*^2^, p = 0.072; fig 5a, d). Turning towards neighbors is not observed in 7 (mean = 0.012 *Rad*⁄*s*^2^, p = 0.301) or 28 dpf surface fish (median = −0.023 *Rad*⁄*s*^2^, p = 0.331; fig 6a & 6d). Similar to trends observed in speeding force, both surface fish and cave fish turn away from fish located within ∼0.1 tank radii across development (fig 5a, b). However, the attractive turning force of cave fish did not significantly differ from zero, except at 28 dpf (7 dpf mean = 0.003 *Rad*⁄*s*^2^, p = 0.919; 28 dpf mean = 0.041 *Rad*⁄*s*^2^, p = 0.0185; 42 dpf mean = 0.048 *Rad*⁄*s*^2^, p = 0.139; 70 dpf mean = −0.045 *Rad*⁄*s*^2^, p = 0.094). Taken together, these findings indicate that surface and cave fish utilize both speeding and turning to maintain preferred positions to neighbors, but only in surface fish during later developmental stages is turning utilized to maintain closer proximities to neighbors. This suggests that the evolved loss of shoaling in cave fish is the product of a loss of these attractive trends in both speeding and turning.

**Figure 5:**
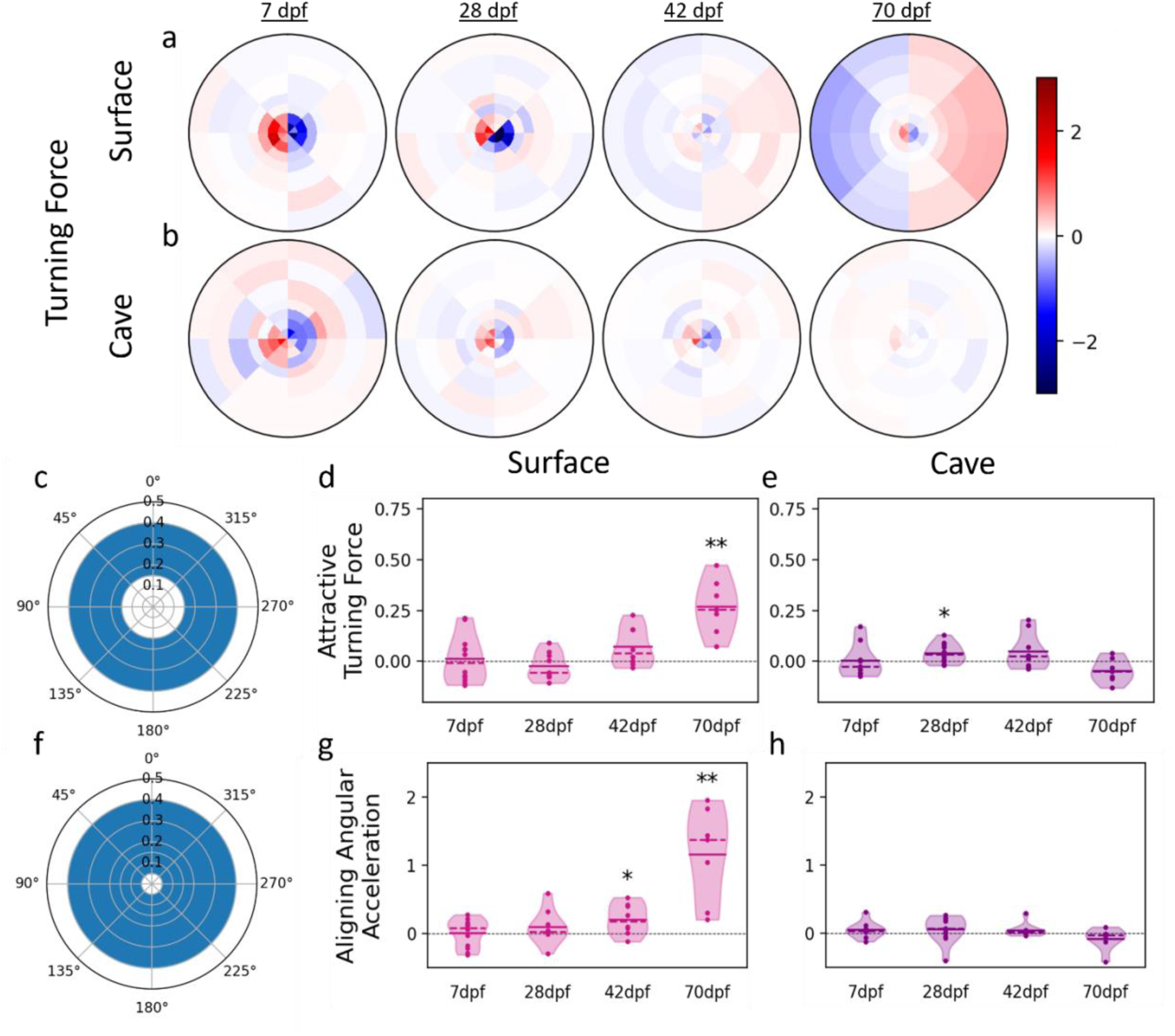
Fish perform turns to control alignment and proximity to neighbors. Turning force represents acceleration perpendicular to a focal **a)** surface or **b)** cave fish’s axis of motion. Turning force here is given as a function of the position of neighboring fish, with the focal fish’s direction of motion oriented toward the top of the figure. Values close to 1 (red) indicate acceleration to the right and values close to −1 (blue) indicate acceleration to the left. **c)** The mean turning force for each trial was calculated for pairs within the attraction zone to get the attractive turning force. Positive values here indicate turning toward a neighbor and negative values indicate turning away. **d)** Mean attractive turning forces of surface fish at 7 dpf (x̃ = 0.012, T = 0.376, p = 0.714), 28 dpf (x̃ = −0.023, T = −1.04, p = 0.331), 42 dpf (x̃ = 0.071, T = 2.12, p = 0.072) and 70 dpf (x̃ = 0.270, T = 5.2, p = 0.002) compared to zero. **e)** Mean attractive turning forces of cave fish at 7 dpf (x̃ = 0.003, T = 0.105, p = 0.919), 28 dpf (x̃ = 0.041, T = 2.81, p = 0.0185), 42 dpf (x̃ = 0.048, T = 1.64, p = 0.139), and 70 dpf (x̃ = −0.045, T = 1.98, p = 0.094) compared to zero**. f)** The contribution of turning to maintaining alignment with neighbors was assessed by comparing the mean trial angular acceleration of pairs within 0.05 and 0.4 tank radii of each other (see methods) **g)** Mean angular accelerations of surface fish at 7 dpf (x̃ = 0.007, T = 0.124, p = 0.904), 28 dpf (x̃ = 0.074, T = 1.16, p = 0.280), 42 dpf (x̃ = 0.210, T = 2.61, p = 0.035), and 70 dpf (x̃ = 1.16, T = 4.45, p = 0.004) compared to zero. **h)** The mean angular acceleration of cave fish did not differ from zero at 7 dpf (x̃ = 0.050, T = 1.25, p = 0.247), 28 dpf (x̃ = 0.061, T = 1.05, p = 0.319), 42 dpf (x̃ = 0.047, T = 6, p =0.055), or 70 dpf (x̃ = 0.023, T = 5, p = 0.156). Accelerations here have been standardized to mean swimming speed of each age and population. * denotes p < 0.05, ** denotes p < 0.01, x̃ = mean, α = 0.05. Results here are the difference between real and mock data.

Fish may utilize turning to alter both their proximity and their alignment relative to other individuals. In order to assess the contribution of turning to the tendency of fish to align with neighbors we calculated the aligning angular acceleration of neighbors located within 0.05 – 0.4 tank radii of each other (see methods; fig 5f). The aligning angular acceleration was calculated as the rate of change of the angular velocity, with a minus sign when the neighbor is on the left of the focal fish so that positive values always correspond to an effort (a torque) to align with the neighbor’s heading whereas negative values indicate an effort to turn away from the neighbor’s heading. At both 42 dpf (mean = 0.210 *Rad*⁄*s*^2^, p = 0.035) and 70 dpf (mean = 1.16 *Rad*⁄*s*^2^, p = 0.004), surface fish angular acceleration is significantly higher than zero, indicating that turning contributes significantly to surface fish tendency to align with neighbors at these ages (fig 5g). In contrast, the angular acceleration of cave fish was close to zero across development, suggesting that fish neither turned to align nor to misalign with neighbors (7 dpf mean = 0.050 *Rad*⁄*s*^2^, p = 0.247; 28 dpf mean = 0.061 *Rad*⁄*s*^2^, p = 0.319; 42 dpf mean = 0.047 *Rad*⁄*s*^2^, p =0.055; 70 dpf mean = 0.023 *Rad*⁄*s*^2^, p = 0.156; fig 5h). Taken together, these results indicate that at late stages of development in surface fish, turning contributes to a preference for being aligned, whereas cave fish have evolved a reduced tendency to turn to align to neighbors.

## Discussion

Collective motion is a complex emergent property that arises from interactions between individuals at the local level (Ariel et al., 2014; Ariel & Ayali, 2015; Bierbach et al., 2020; Corcoran & Hedrick, 2019; Herbert-Read et al., 2011, 2017; Katz et al., 2011; Knebel et al., 2019; Young et al., 2013). Schooling and shoaling in fish are examples of collective motion, and while the local interactions that underlie schooling and shoaling have been studied in several fish species (Bierbach et al., 2020; Herbert-Read et al., 2011; Katz et al., 2011), little is known about how natural variation in these local interactions result in evolved differences at the level of the emergent collective behavior (Greenwood et al., 2013). There is considerable variation in schooling and shoaling among fish that live in different ecological conditions. For example, populations of Trinidadian guppies display different degrees of group cohesion, and cohesiveness positively correlates with the degree of predation in their natural habitats (Herbert-Read et al., 2017; Huizinga et al., 2009; Ioannou et al., 2017; Magurran et al., 1992; Seghers, 1974; Song et al., 2011). Additionally, sociality of threespine stickleback populations varies according to water temperature during rearing – a trend that is particularly worrisome as global temperatures rise (Pilakouta et al., 2023). While there is significant diversity in the tendency to school and shoal across populations of fishes, how evolution impacts local interaction rules to produce these group level differences is not understood. Here we illustrate how evolved changes in interindividual interactions culminate in the development of naturally occurring differences in schooling and shoaling in closely related populations of a single species.

While studies in zebrafish have laid the groundwork for understanding how complex collective behaviors manifest over the course of development, *Astyanax mexicanus* represents a unique opportunity to not only determine how collective behaviors emerge over development, but also how the individual behaviors and pairwise interactions that underlie collective motion change when emergent behaviors evolve. Observations of differences in the collective behaviors of adult surface and cave populations of the Mexican tetra go back at least as far as 1964 (John, 1964), and include both field and lab studies (Gregson & Burt de Perera, 2007; John, 1964; Kowalko et al., 2013; Patch et al., 2022). However, it was only recently that studies began capitalizing on automated tracking, allowing for previously unattainable in-depth quantitative analysis of these behaviors (Iwashita & Yoshizawa, 2021; Patch et al., 2022). These studies also demonstrate that, although they do not exhibit robust schooling and shoaling, adult cave fish modulate their behavior in the presence of conspecifics by altering average swimming and turning speeds, and that their sociality may be altered by environmental conditions (Iwashita & Yoshizawa, 2021; Patch et al., 2022). Thus, cave fish present an opportunity to understand mechanisms contributing to evolution of collective behaviors. Here, we assess differences in how individual fish respond to other individuals in these populations of schooling and non-schooling fish. We find that by 70 dpf, the collective behaviors of surface and cave fish resemble those of adults of the same populations assayed under similar conditions (Patch et al., 2022), with surface fish exhibiting robust schooling and cave fish displaying no attraction or tendency to align. Further, we have characterized the individual level behavioral changes that underlie these differences in schooling and shoaling: Surface fish utilize turning and changes in speed to maintain close proximities and alignment relative to neighbors, similar to individuals from other schooling species (Harpaz et al., 2021; Herbert-Read et al., 2011; Katz et al., 2011). Changes in swimming speed are utilized by surface fish to control proximity to neighbors, while turning is utilized to control both proximity and alignment to neighbors. In contrast, while cave fish also utilize turning and speed changes to maintain control their positions relative to neighbors, they do not utilize turning to alter their alignment relative to neighbors. Furthermore, cave fish only perform turns and modulate speed to separate from neighbors, they do not perform turns or modulate speed to maintain close proximities to neighbors, resulting in the loss of the robust attraction and alignment found in schooling and shoaling fish. Similarly, experiments in which groups of guppies were artificially selected for greater group alignment across multiple generations also resulted in changes in the relationship between both turning and changes in speed relative to the positions of neighbors (Kotrschal et al., 2020). Groups selected for greater cohesion exhibited stronger correlations between turning and nearest neighbor direction. The correlation between turning and nearest neighbor position in groups selected for cohesion was not significantly stronger than in non-selected groups though a slight trend was observed (Kotrschal et al., 2020). These findings, along with our results in *A. mexicanus*, support the idea that the modulation of turning and speed is essential to evolved differences in collective behavior across fish species.

One approach for understanding how changes at the level of local interactions affect schooling and shoaling is to perform in-depth quantitative analyses of the interactions across developmental timepoints. To the best of our knowledge, analyses of the development of schooling and shoaling had exclusively been conducted in the zebrafish, *Danio rerio*, prior to this study (Harpaz et al., 2021; Hinz & de Polavieja, 2017; Stednitz & Washbourne, 2020). Similar to our findings in the Mexican tetra, zebrafish develop attraction and tendency to align to other fish at distinct points in development (Harpaz et al., 2021; Hinz & de Polavieja, 2017; Stednitz & Washbourne, 2020). In accordance with these findings, previous genetic screens have indicated that these components of schooling and shoaling are regulated by different genes (W. Tang et al., 2020). Unlike in the Mexican tetra, attraction precedes the tendency to align in zebrafish larvae, and both attraction and tendency to align emerge much earlier in development in zebrafish (Harpaz et al., 2021; Stednitz & Washbourne, 2020). Similar to our results in surface *A. mexicanus*, developmental changes in tendency to align and attraction in zebrafish are the product of changes in the ways individual fish modify their velocity in response to neighbors (Harpaz et al., 2021). Whether the differences in ontogeny of schooling across species are due to life history or differences in ecological factors under which these populations have evolved remains an open question.

Analysis of ontogeny of schooling and shoaling in cave and surface fish revealed similarities between the inter-individual interactions of both populations early in development. In addition to similar trends in nearest neighbor pair angle, interindividual distance, and nearest neighbor distance when compared to control mock data at early stages of development, analysis of individual behaviors relative to neighbors reveals that both cave fish and surface fish modulate speed and turning to create distance between themselves and close neighbors, and do not modulate speeding or turning in response to more distant neighbors at early developmental stages, including those in the area that makes up the attraction zone at later stages of development in surface fish. Beginning around 42 dpf, however, the developmental trajectories of these behaviors diverge between surface and cave fish. Surface fish begin to develop robust attractive interactions, turning toward neighbors and modulating speed to remain close to neighbors, as well as aligning interactions, turning to align with neighbors. Neither attraction nor alignment are observed in cavefish, which do not turn nor speed to get closer to or align with neighbors. These data suggest that the evolution of collective behaviors is driven by changes in patterns of speeding and turning by individuals to move closer to or align with neighbors.

Intriguingly, differences in social interactions between surface fish and cave fish are not present at early developmental stages, as fish which in later development do not school and shoal still modulate turning and speeding in response to close neighbors, suggesting a model for how schooling and shoaling evolve: through a loss of behaviors that result in attraction and alignment, rather than loss in all modulation of behavior based on location of other fish, or through cave fish exhibiting school-promoting interactions, but which are too weak to yield actual schooling (as is suggested by analysis of adult surface fish in the dark (Patch et al., 2022)). These results, combined with recent advances in *A. mexicanus* research, including Tol2 transgenesis (Stahl et al., 2019), CRISPR gene editing (Klaassen et al., 2018), and neuroanatomical brain atlases (Jaggard et al., 2020; Kozol et al., 2022; Loomis et al., 2019), will provide a unique opportunity to probe for the neuronal and genetic mechanisms underlying naturally occurring variation in components of collective behavior in future studies.

Collective behaviors are exhibited by a wide variety of animals and, like schooling and shoaling in fishes, are the product of local interactions between individuals (Ariel et al., 2014; Ariel & Ayali, 2015; Bierbach et al., 2020; Corcoran & Hedrick, 2019; Herbert-Read et al., 2011, 2017; Knebel et al., 2019; Young et al., 2013). Indeed, this trend applies not only to animals but also to groups of cells or even moving particles (Barriga & Mayor, 2015; Bhattacharjee et al., 2022; Czirók & Vicsek, 2000; Eglinton et al., 2022; Theveneau & Mayor, 2013). The ability then to understand how changes in local interactions influence collective behaviors is relevant to a wider variety of disciplines than simply animal behavior, emphasizing the significance of data such as those presented here.

## Materials and Methods

### Animal care

Surface and cave embryos were collected the morning after spawning and placed in glass Pyrex bowls filled with conditioned fish water. >1 dpf embryos were sorted into groups of 50 in 350 ml glass Pyrex bowls. After being assayed at 7 dpf, fish were transferred into 2L plastic tanks where they remained until 14 dpf. Fish were then transferred into 6L tanks on a filtered aquatic housing system, where they remained for the rest of the experiment. Prior to being placed on the system, routine water changes were performed. Beginning at 6 dpf, fish were fed twice a day on weekdays and once a day on weekends. All individuals received a combination of brine shrimp and GEMMA Micro. All cave fish used in these assays were descendants of adult fish originally collected from the Pachón cave, and all surface fish used were descendants of individuals originally collected from rivers in Mexico and Texas. All protocols were approved by the IACUC of Florida Atlantic University, and all fish were kept in Florida Atlantic University fish facilities. Water temperatures were maintained at 23 ± 1°C and light:dark cycles were kept at 14:10, with a light intensity between 24 and 40 lux.

### Behavioral experiments

All fish were fed to satiety at least 1 hour before beginning assays. Before being assayed, fish were carefully netted into a holding tank for one minute and then were gently poured into a circular arena and allowed to acclimate for 10 minutes. After the acclimation period, behavior was recorded for a duration of 20 minutes at 30 fps with a video camera (FLIR; GS3-U3-23S6M-C) equipped with a wide-angle c-mount lens (Edmund Optics; HP Series 12 mm fixed focal length lens) mounted above the center of the arena on a custom stand constructed from polyvinyl chloride (PVC) tubing. Assays were recorded as series of .RAW files which included timestamps for each frame. Arena diameters were increased across developmental timepoints assayed in order to maintain approximately a ratio of 22 body lengths per arena diameter (table 1; fig S4a & b)). Arenas were 3D printed (Creality; CR10MAX) in black polylactic acid (PLA) and adhered onto a sheet of clear acrylic with acrylic cement and then rendered waterproof with a layer of silicone along the base of the outer edge of the arena. Arenas were placed on top of custom-made white acrylic boxes (76 x 76x 14 cm) that diffused light emitted by white-light LED strips placed under the box.

**Table 1.** Arena specifications across time points. Arena diameters and depths, in mm, for each

| Age | <u>7 dpf</u> | <u>28 dpf</u> | <u>42 dpf</u> | <u>70 dpf</u> |
| --- | --- | --- | --- | --- |
| Arena<br>(diameter x<br>depth)<br>(mm) | 96 x 7 | 177 x 7 | 242 x 25 | 339 x 25 |

### Tracking

RAW files were compiled into videos (.mkvs) and subsequently processed using version 0.1.1 of the custom python tracking library trilab-tracker (Patch et al., 2022) (located at https://github.com/yffily/trilab-tracker/releases/tag/0.1.1), which extracts the positions and orientations of fish. All tracking and orientation data were manually verified, and corrections were applied when necessary. Arena edges were selected manually and used to convert pixels to centimeters based on the arena diameter. Trajectories were smoothed using a five-frame Savitzky-Golay filter (scipy.signal.savgol_filter with window_length=5). Fish velocities and accelerations and their angular counterparts were computed using standard finite difference formulas:

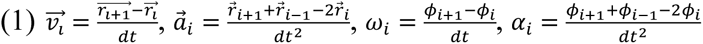

Where 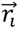 is the fish’s position vector in frame number *i*, 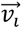 is its velocity vector, 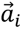 is its acceleration vector, 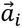 is the angle between the x axis and the fish’s orientation, *w*_*i*_ is the fish’s angular velocity, and α_*i*_ is the fish’s angular acceleration.

### Mock group formation

All possible combinations of five trials were found for each combination of age and population. For each combination of five trials, the tracks for a random fish were chosen from each trial and combined to form a mock group, so that each mock group contained five fish and the quantity of mock groups was equal to

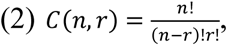

where n is the quantity of real trials and r = 5 (the number of fish per trial). Because fish are randomly chosen for each group, multiple iterations of the mock group formation process will result in different outcomes. In order to assess variability across iterations, the outcomes of 10 iterations were compared via Kruskal-Wallis and found to be similar. The results of the first iteration were used for all comparisons between mock groups and real groups.

### Orientation and distance analyses

Pair distance was calculated by finding the distance between the center points of two fish. Interindividual distance was calculated by finding the pair distance between a focal fish and all other fish in the trial and nearest neighbor distance was calculated by finding the minimum pair distance for each fish in each frame. Pair angle was calculated by finding the difference between the orientations of two fish. Nearest neighbor pair angle was calculated by finding the pair angle between the focal fish and its nearest neighbor. Nearest neighbor pair orientations, nearest neighbor distances, and interindividual distances were measured for real and mock data. Before hypothesis testing, distributions of distance and pair angle data were assessed using Shapiro-Wilk tests for real data and Kolmogorov-Smirnov tests for mock data to account for differences in quantities of real and mock groups. Comparisons were made between real and mock data of the same age and population using a student’s t-test if both the real and mock data were found to be normally distributed or a Mann-Whitney U test if either dataset was found to be non-normal. Throughout the text, means are reported for data that were found to be normally distributed and medians are reported for data that were not found to be normally distributed. As fish can develop at different rates, we also assessed the relationship between these metrics and body length. While analyses herein were conducted according to the age of the fish, body length is also a good predictor of changes in proximity (fig S5). The correlations between swimming speed and proximity and alignment were assessed by calculating the Spearman’s rank correlation coefficient for each age and population using the mean trial swimming speed and mean nearest neighbor distance, mean interindividual distance, or mean nearest neighbor pair angle for each frame of each trial.

### Density heatmaps

Density heatmaps (fig 3) show the density of fish around a focal fish located at the center of the heatmap, facing up. First a focal fish is picked. The focal fish’s coordinate system is defined, whose origin is the center point of the focal fish and whose y axis points in the direction faced by the focal fish. The coordinates of every other fish in the trial are computed in this coordinate system, normalized by the tank radius, then binned according to figure 3. The radius bin edges are 0, 0.05, 0.1, 0.15, 0.2, 0.3, 0.4, and 0.5. The value of each bin is the probability of finding a fish in that bin, divided by the area of that bin. The result is in fish per square tank radii.

### Force heatmaps

Force heatmaps (fig 4, 5) show the average speeding and turning forces of the focal fish when another fish is present nearby as a function of the location of that other fish. The speeding force is the component of the focal fish’s acceleration that is parallel to its own orientation, i.e., the focal fish’s tangential acceleration. The turning force is the component of the focal fish’s acceleration that is perpendicular to its own orientation, i.e., the focal fish’s normal acceleration. The latter is counted positively if it points to the right of the fish and negatively to the left. The acceleration is computed using finite differences. The orientation is obtained from the fish’s body shape. Both accelerations are normalized by the average swimming speed of the fish’s population and age cohort. Once a focal fish has been picked, the coordinates of the other fish in the trial are computed and binned as for density maps. The value of a bin is the average of the focal fish’s speeding or turning force over every frame in which there was a second fish in that bin. Overall, our method is similar to the one used by Katz et al. (Katz et al., 2011) except our bins are based on polar rather than cartesian coordinates and they do not overlap.

### Attractive speeding and turning force

The speeding force is positive when the focal fish speeds up and negative when it slows down. If the focal fish uses speed changes to get closer to its neighbors, we expect it to speed up when the neighbor is ahead but slow down when the neighbor is behind. Conversely, speeding up when the neighbor is behind and slowing down when the neighbor is ahead suggests repulsion. Therefore, we define the attractive speeding force to be equal to the speeding force when the neighbor is ahead but minus the speeding force when the neighbor is behind. With this definition, positive values indicate attraction and negative values indicate repulsion. Similarly, we define the attractive turning force to be equal to the turning force when the neighbor is on the right side of the focal fish but minus the turning force when the neighbor is on the left side of the focal fish. With this definition, positive values indicate attraction (the focal fish’s trajectory is curving towards the neighbor) and negative values indicate repulsion (the focal fish’s trajectory is curving away from the neighbor). The attractive speeding and turning forces are then averaged over all possible locations of the neighbor fish, restricted to the range of distances where we expect attraction. The density heatmap for 70dpf surface fish, which exhibit robust schooling, shows a ring of increased probability between about 0.05 and 0.4 tank radii around the focal fish, with a peak around 0.15 tank radii. Therefore, we expect interactions to be repulsive on average between 0.05 and 0.15 tank radii and attractive on average between 0.15 and 0.4 tank radii. Short range repulsion may be simple collision avoidance, so we focus on the attractive range, i.e., distances between 0.15 and 0.4 tank radii. The average over neighbor positions is weighted by each bin’s area, i.e., all possible location of the neighbor fish within the allowed distance range are treated equally, independently of the likeliness of finding a fish there. Weighing instead by the likeliness of finding a fish in each bin yields similar results. Speeding and turning force violin plots are the difference between real and mock data (fig S6a-d).

### Aligning angular acceleration

Just like the attractive speeding and turning forces are defined to be positive when they contribute to decreasing the distance to the focal fish’s neighbor, the aligning angular acceleration is defined to be positive when it contributes to decreasing the angle between the headings of the focal fish and its neighbor. We start with the angular acceleration, which is positive when the focal fish attempts to rotate counterclockwise and negative when it attempts to rotate clockwise, then flip the sign if the angle between the heading of the focal fish and the heading of the neighbor fish is negative (between −180° and 0°). We then average over neighbor locations whose distance to the focal fish is between 0.05 and 0.4 tank radii. The upper bound (0.4 tank radii) is the same use to compute the average attractive speeding and turning forces. The lower bound (0.05 tank radii) is lower than the one used for the average attractive speeding and turning forces (0.15 tank radii) because while we expect schooling fish less than 0.15 tank radii away from each other to attempt to maintain alignment while they adjust their distance to each other. Angular acceleration violin plots are the difference between real and mock data (fig 6e & f).

### Density, force, and angular acceleration mock data

The mock data used in figures 4 to 6 were obtained by averaging the relevant quantity (speeding force, turning force, or angular acceleration) over every possible pair of fish taken from two different trials. This is equivalent to averaging over every pair of fish from the same mock trial and every possible mock trial (every possible group of 5 fish taken from 5 different real trials). This only works because all quantities shown figures 4 to 6 are pairwise. It would not work for, e.g., a quantity involving the nearest neighbor as the identity of the nearest neighbor depends on the position of every fish in the trial.

### Statistical analysis of attractive forces and aligning angular acceleration

After subtracting the mean mock value, the distribution of attractive speeding force, attractive turning force, or aligning angular acceleration was tested for normality using a Shapiro-Wilk test, then compared with zero using either a one-sample T-test (if the data was normal) or a Wilcoxon test (if the data was not normal).

### Analysis software

Figures were generated and analyses performed using custom Python 3 scripts that will be share upon request. The Pandas and Numpy libraries were used for data organization and analysis. Statistical analysis was performed using the following python libraries: scipy.stats for Kolmogorov-Smirnov, Shapiro-Wilk, Kruskal-Wallis, Mann-Whitney U, Wilcoxon Signed-Rank test, and t-tests; and scikit_posthocs for Dunn’s test. Figures were generated using the Matplotlib and Seaborn libraries.

## Acknowledgments

We would like to thank Adam Patch for advice with data analysis. This study was supported by National Institutes of Health grant R35GM138345 to JEK, National Science Foundation grant IOS2202359 to JEK and National Science Foundation EDGE grant 1923372 to JEK and ERD.

## Data and materials availability

Tracking software is available at https://github.com/yffily/trilab-tracker/releases/tag/0.1.1

## Author contributions

Conceptualization: AP, JEK, YF; Methodology: AP, JEK, YF; Investigation: AP, KJH, AC, AA, BA, JEK; Visualization: AP, YF; Supervision: JEK, YF, ERD, ACK; Writing—original draft: AP, JEK; Writing—review & editing: AP, JEK, YF, ERD, ACK, KJH, AC, AA, BA

