## Supplemental Files for "Changes in local interaction rules during ontogeny underlie the evolution of collective behavior"

Surface

| a | | Nearest Neighbor ϴ° | Nearest Neighbor Distance | Interindividual Distance |
|---|---|---|---|---|
| 7 dpf | $r_s$ | 0.02 | -0.0474 | -0.0314 |
| | $R^2$ | 0.000 | 0.002 | 0.001 |
| | p | $p < 0.001$ | $p < 0.001$ | $p < 0.001$ |
| 28 dpf | $r_s$ | -0.009 | -0.016 | -0.008 |
| | $R^2$ | 0.000 | 0.000 | 0.000 |
| | p | $p < 0.001$ | $p < 0.001$ | $p < 0.001$ |
| 42 dpf | $r_s$ | -0.162 | -0.139 | -0.228 |
| | $R^2$ | 0.026 | 0.019 | 0.052 |
| | p | $p < 0.001$ | $p < 0.001$ | $p < 0.001$ |
| 70 dpf | $r_s$ | -0.185 | 0.141 | 0.205 |
| | $R^2$ | 0.034 | 0.020 | 0.042 |
| | p | $p < 0.001$ | $p < 0.001$ | $p < 0.001$ |

Cave

| b | | Nearest Neighbor ϴ° | Nearest Neighbor Distance | Interindividual Distance |
|---|---|---|---|---|
| 7 dpf | $r_s$ | -0.001 | 0.018 | 0.081 |
| | $R^2$ | 0.000 | 0.004 | 0.026 |
| | p | 0.005 | $p < 0.001$ | $p < 0.001$ |
| 28 dpf | $r_s$ | -0.001 | 0.018 | 0.081 |
| | $R^2$ | 0.000 | 0.000 | 0.006 |
| | p | $p < 0.001$ | $p < 0.001$ | $p < 0.001$ |
| 42 dpf | $r_s$ | 0.021 | 0.017 | 0.177 |
| | $R^2$ | 0.000 | 0.000 | 0.031 |
| | p | $p < 0.001$ | $p < 0.001$ | $p < 0.001$ |
| 70 dpf | $r_s$ | 0.057 | 0.199 | 0.436 |
| | $R^2$ | 0.003 | 0.040 | 0.190 |
| | p | $p < 0.001$ | $p < 0.001$ | $p < 0.001$ |

**Fig S1. Correlations between swimming speed and proximity and alignment.** Spearman's rank correlation coefficients were calculated in order to assess correlations between swimming speed and nearest neighbor pair angle, nearest neighbor distance, and interindividual distance for **a)** surface and **b)** cave fish. $r_s$ denotes Spearman's rho, $R^2$ denote R-squared, and p denotes to p-value for each comparison.

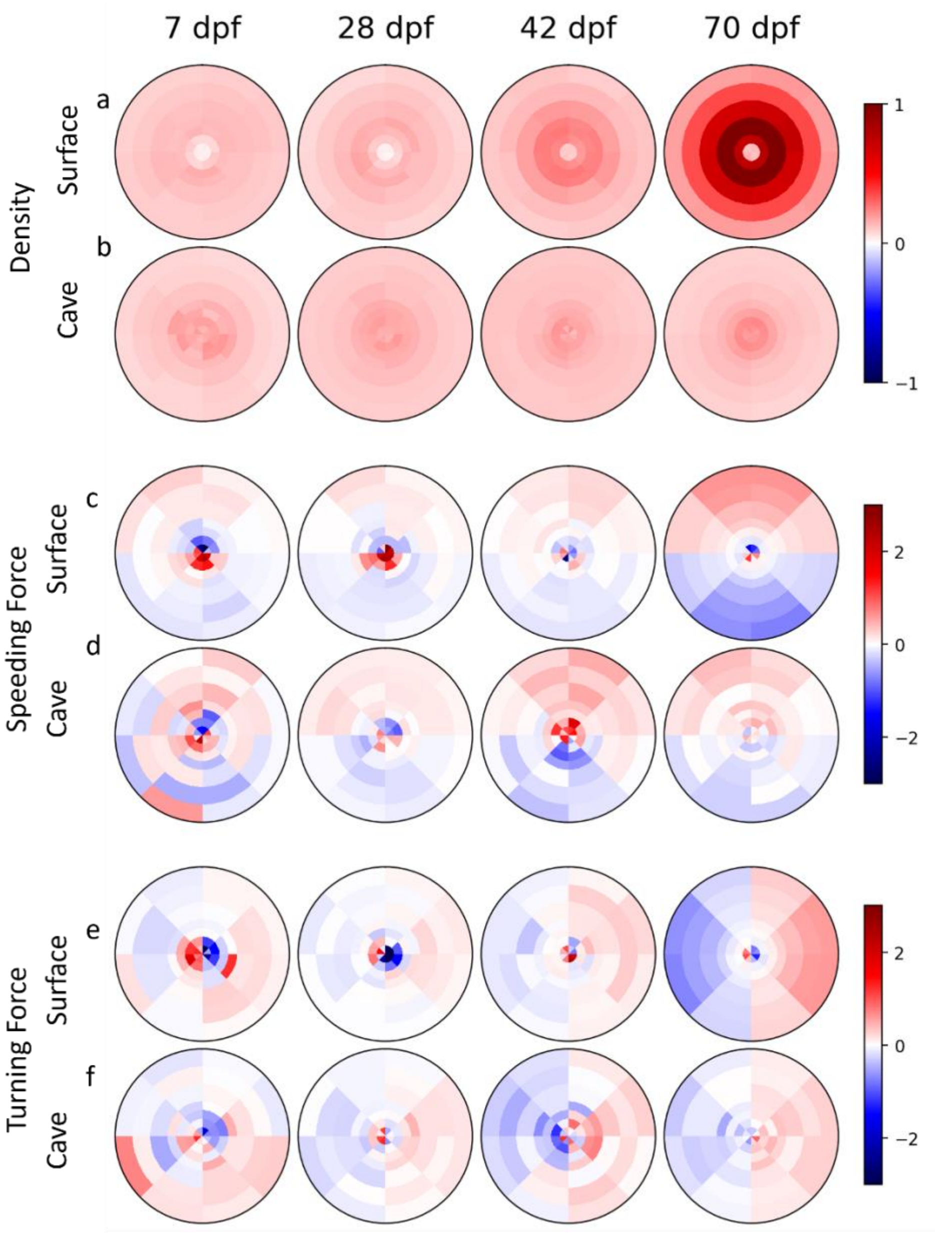


**Fig S2. Density, speeding force, and turning force of groups of five fish.** Density heat maps showing positional preferences of real groups of **a)** surface and **b)** cave fish. Heat maps illustrating speeding force of real groups of **c)** surface and **d)** cave fish. Heat maps illustrating the turning force of real groups of **e)** surface and **f)** cave fish.

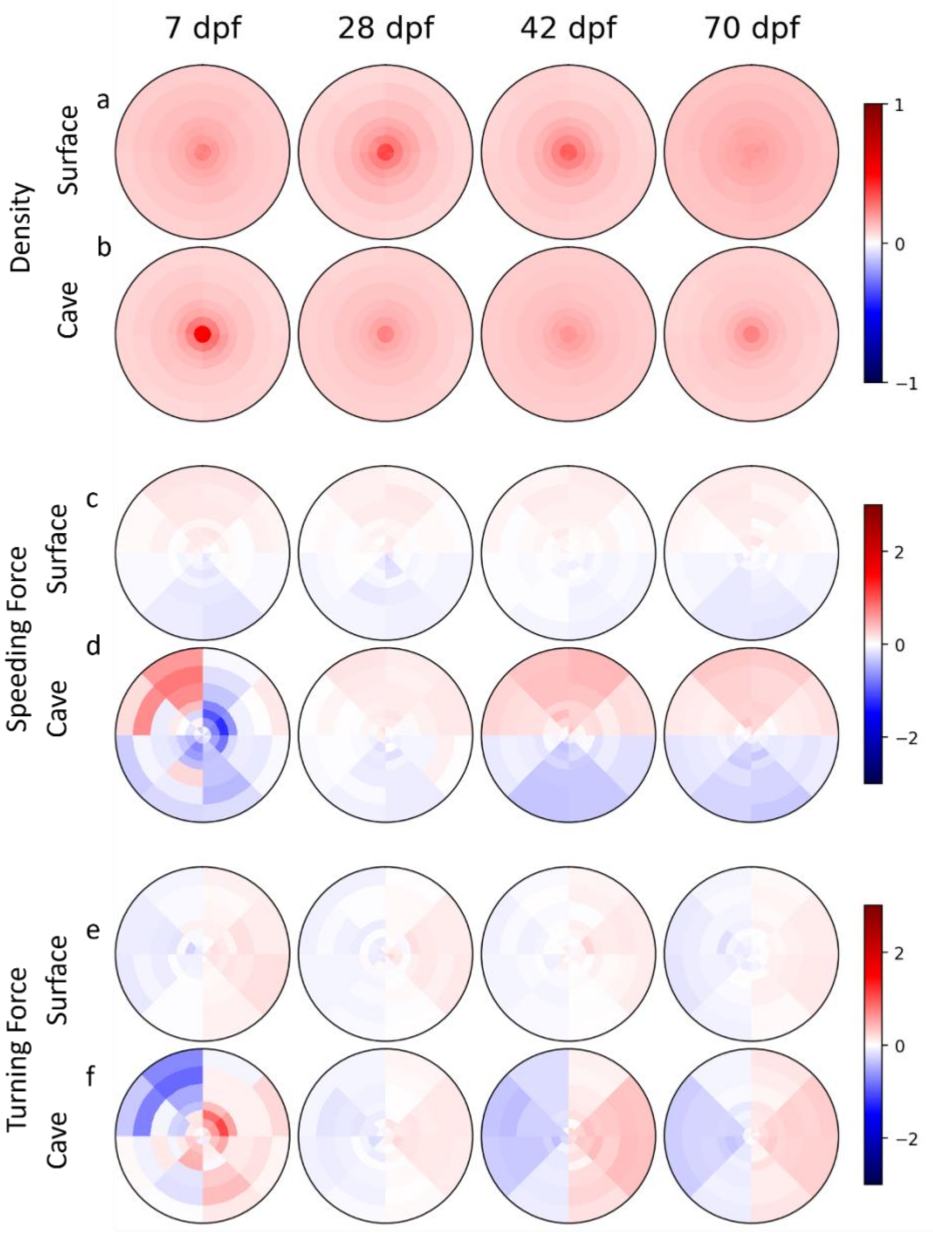


**Fig S3. Density, speeding force, and turning force of mock groups of five fish.** Density heat maps showing positional preferences of mock groups of **a)** surface and **b)** cave fish. Heat maps illustrating speeding force of mock groups of **c)** surface and **d)** cave fish. Heat maps illustrating the turning force of mock groups of **e)** surface and **f)** cave fish.

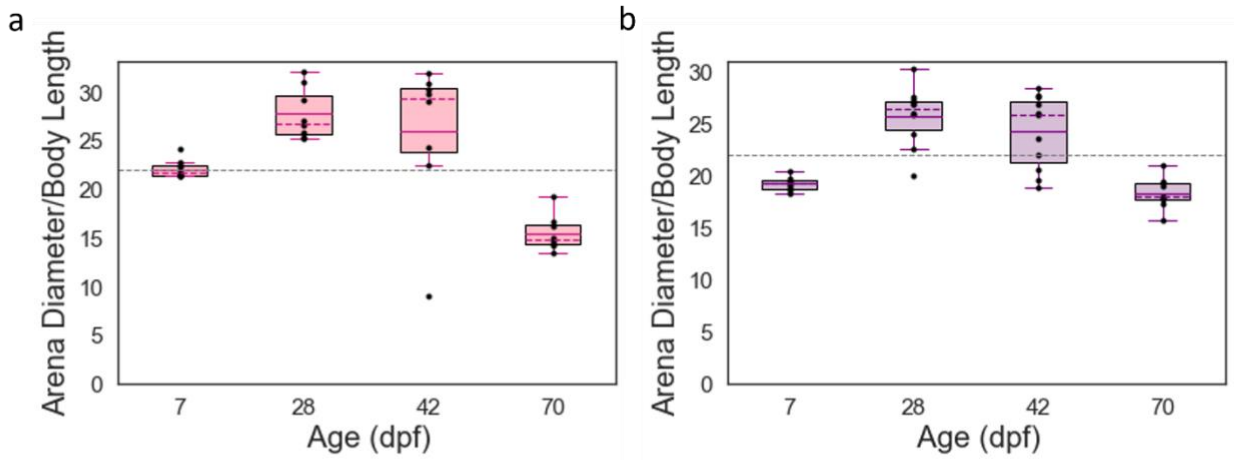


**Fig S4. Ratio of fish body length to arena diameter.** Ratio of body length to arena diameter for **a)** surface fish and **b)** cave fish. Solid line within each boxplot denotes mean and dashed line within each boxplot denotes median. Each point denotes the mean body length for a single trial with of fish. Dashed lines across figures denote desired 1 diameter:22 body lengths ratio.

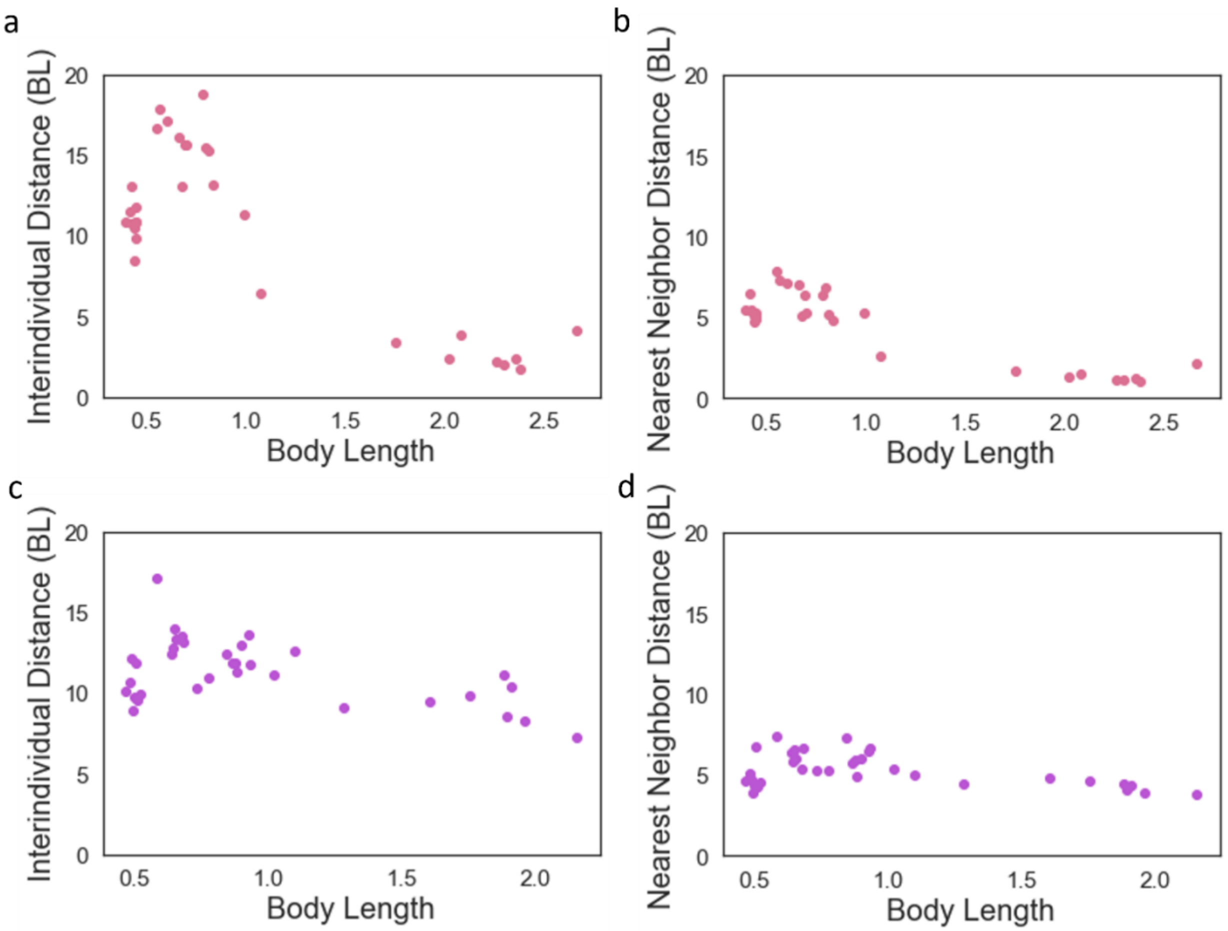


**Fig S5. Associations between proximity and body length.** Surface fish **a)** interindividual and **b)** nearest neighbor distances standardized to body length and given relative to body length. Cave fish **c)** interindividual and **d)** nearest neighbor distance standardized to body length and given relative to body length. Each point represents the average body length and median nearest neighbor or interindividual distance per trial. Purple = cave fish, purple = cave fish.

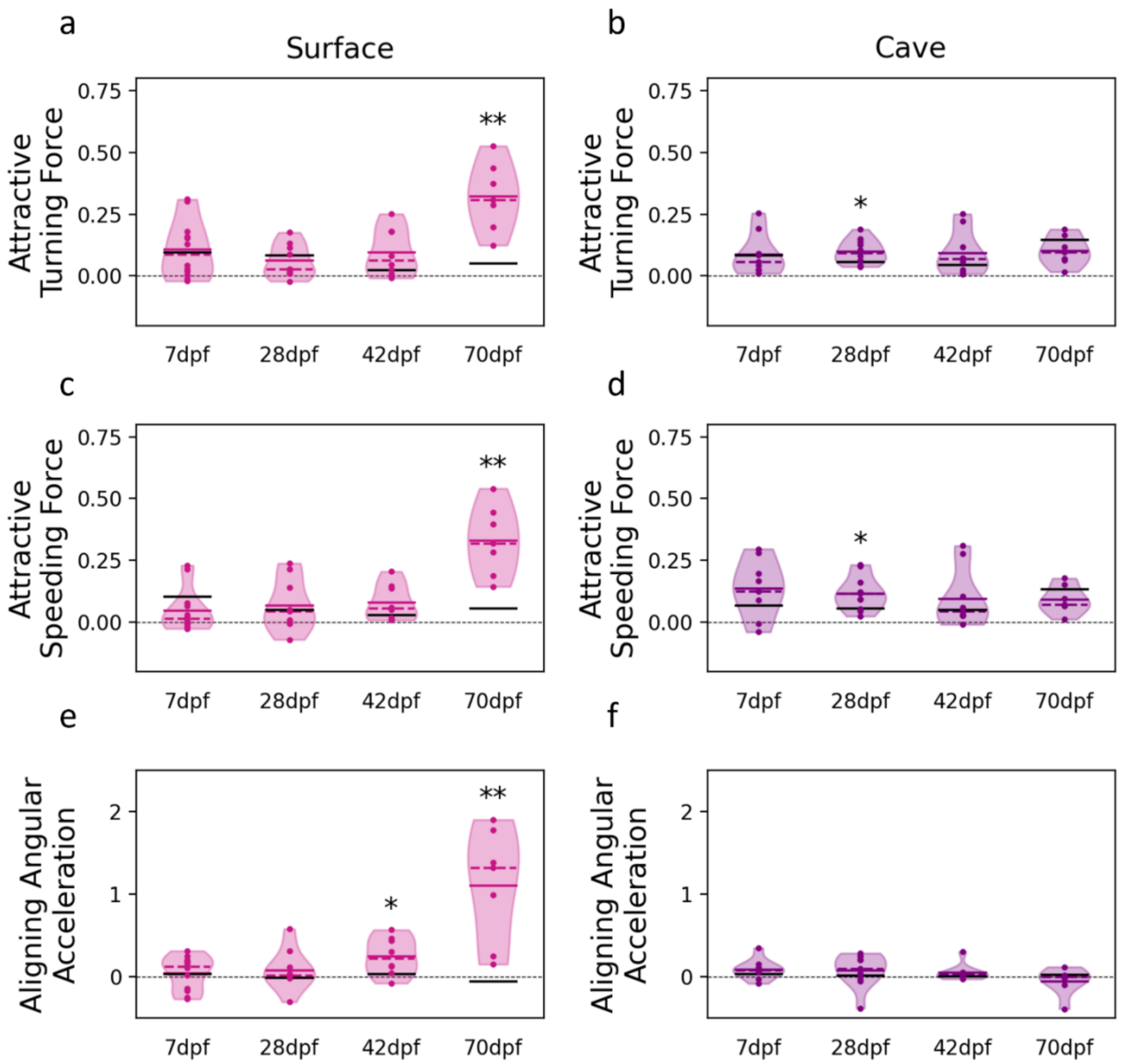


**Fig S6. Real and mock attractive turning and speeding force and aligning angular acceleration.** Real and mock attractive turning force for surface (a) and cave (b) fish. Attractive speeding force for surface (c) and (d) cave fish. Aligning angular acceleration for surface (e) and cave (f) fish. Real surface fish data in pink, real cave fish data in purple, and mean mock value as a black bar. Solid lines denote means, dotted lines denote medians.
